# Universality in biodiversity patterns: Variation in species-temperature and species-productivity relationships reveals a prominent role of productivity in diversity gradients

**DOI:** 10.1101/2020.12.16.422752

**Authors:** Eliška Bohdalková, Anna Toszogyova, Irena Šímová, David Storch

## Abstract

Temperature and productivity appear as universal positive correlates of species richness. However, the strength and the shape of species-temperature (STR) and species-productivity (SPR) relationships vary widely, and the causes of this variation are poorly known. We analysed (1) published species richness data for multiple taxa sampled in various regions and (2) different clades within vertebrate classes globally, to test for the effects of spatial scale and characteristics of examined taxa and regions on the strength and direction of STRs and SPRs. There are striking differences in the variation of the relationships among types of data, between ectotherms and endotherms and also between STRs and SPRs. Some sources of this variation are of statistical nature (e.g. the relationships are stronger if the range of temperature or productivity variation is wider), but non-statistical sources are more important and illuminate the processes responsible for the origin of biodiversity patterns. The SPRs are generally stronger and less variable than STRs, and SPR variation is weakly related to the explored factors – the SPRs are stronger in warmer regions in ectotherms, while clade size is the only factor consistently affecting the strength of the SPR in endotherms. In contrast, STRs are weaker and more variable, and this variation is linked to region characteristics - most importantly, STRs are stronger in the regions where temperature positively correlates with productivity, indicating that productivity plays a role even in the STRs. The effect of temperature on species richness is thus complex and context-dependent, while productivity is a more universal driver of species richness patterns, largely independent of particular characteristics of given region or taxon. Productivity thus appears as the main proximate driver of species richness patterns, probably due to its effect on the limits of the number of viable populations which can coexist in a given environment.

## Introduction

Deciphering large-scale diversity patterns is one of the main goals of ecology (Gaston 2000, Pontarp et al. 2019). It has been shown that species richness is tightly related to climate, namely to temperature, water availability and resulting ecosystem productivity (Wright 1983, Currie 1991, Hawkins et al. 2003a, Field et al. 2009), so that temperature and productivity appear as the main variables positively affecting species richness (Lennon et al. 2000, Kaspari et al. 2003, Currie et al. 2004, Evans et al. 2005, Šímová et al. 2011). However, the documented species-temperature (STR) and species-productivity (SPR) relationships vary in terms of their strength and shape. Weak or variable STRs or SPRs are sometimes interpreted as a counterevidence against the hypotheses explaining diversity patterns by the processes driven by temperature or productivity (Hawkins et al. 2007, Buckley et al. 2010). Recovering the causes of the variation of STRs and SPRs is thus a necessary precondition for understanding the role of temperature and productivity in diversity patterns.

Productivity and/or temperature are comprised in most of the hypotheses that explain large-scale diversity patterns (Pontarp et al. 2019). Productivity may determine the total amount of resources that can be divided among species to allow the persistence of their viable populations - according to the more-individuals hypothesis, low productivity cannot support high number of species as these would have so small populations that their extinction rates would be higher than origination rates (Storch et al. 2018). Higher productivity may also allow higher species’ specialization promoting coexistence (Abrams 1995). Alternatively, productivity may promote larger population sizes, positively affecting mutation accumulation and speciation (Stevens et al. 2007). Temperature may affect mutation and consequently speciation rates as well (Rohde 1978, Allen et al. 2002) or it may constrain the number of species which can persist in a given environment due to their physiological temperature tolerances (Šímová et al. 2011). Alternatively, the effect of temperature may be indirect via productivity (Šímová and Storch 2017). Higher species richness in warmer and more productive areas may also be a consequence of historically larger and more stable tropical regions (Wiens and Donoghue 2004).

Importantly, synthetic macroecological theories that attempt to explain large-scale diversity patterns via a combination of several fundamental processes typically assume that productivity and temperature represent key drivers of these processes. Specifically, the neutral theory (Hubbell 2001) assumes that species richness within a region is determined by the product of total number of individuals, *J*, and speciation rate, *v*. It is reasonable to assume that *J* is a function of productivity (together with area) of the region, while *v* may be linked to temperature. These assumptions provide the basis of the theories which combine the neutral theory with the metabolic theory of ecology (Allen et al. 2007, Worm and Tittensor 2018), and are also in accord with the equilibrium theory of biodiversity dynamics (Storch et al. 2018, Storch and Okie 2019). Positive effects of productivity and temperature on species richness are thus essential features of major macroecological integrative theories.

On the other hand, there are good theoretical reasons to expect that neither STR nor SPR are universal, and that considerable variation exists around canonical, monotonically increasing relationships. Some of these reasons are statistical – smaller groups of species or small regions which do not comprise enough variation of temperature or productivity may reveal relatively weak and/or insignificant patterns even if there is an overall positive trend of increasing species richness with temperature or productivity. Other reasons may relate to biological specificities of the explored taxa. Many clades would reveal diversity patterns that reflect the spreading from an area of origin rather than environmental effects (Gehrke and Linder 2011). This is especially the case of young (and thus small) clades which did not yet have time to reach an equilibrium, and their species richness patterns are thus driven by the history of their radiation and dispersal. The effects of productivity or temperature on species richness may be also weak if other factors contribute to species richness variation. For example, productivity may be less relevant for taxa which utilize specific resources whose levels do not correlate with net primary productivity. Also, any positive effects of temperature on species richness may be irrelevant when the lack of water limits the very existence of life, e.g. on deserts.

Importantly, the variation of the STR and the SPR may reflect the exact ways that these variables affect species richness. For instance, if temperature affects species richness only indirectly, through its effect on productivity, then the SPR should be more universal (i.e. more regular and context-independent) than the STR, and the effect of temperature should diminish when accounting for productivity. Also, if temperature affects diversity via affecting mutation and speciation rates, the metabolic theory (Brown et al. 2004) predicts exponential STR with a predictable slope. However, this prediction holds only if total community abundance is more or less constant (Allen et al. 2007), while the STR is expected to vary if this condition is not fulfilled (Storch 2012). Since the total number of individuals is constrained by productivity (Storch et al. 2018), productivity variation may, according to the metabolic theory, also affect the slope and strength of the STR. More generally, if both productivity and temperature simultaneously affect species richness, both the STR and the SPR may depend on the variation of the other variable.

The variation of STRs and SPRs has been only rarely studied in a comprehensive way. Several studies addressed the effect of spatial scale and its two components – grain and extent – on biodiversity patterns including the STR and the SPR. Generally, with increasing *grain* size, the relationships are stronger (Rahbek and Graves 2000, 2001, González-Taboada et al. 2007, Belmaker and Jetz 2011), their slope is steeper (Hillebrand 2004), and the relative importance of temperature and productivity changes (Field et al. 2009, Belmaker and Jetz 2011). The effect of geographical *extent* is mostly studied using meta-analyses – and again, with increasing the extent of the analysis, the relationships appear stronger and steeper (Hillebrand 2004) and the importance of climate and productivity increases (Field et al. 2009). Some other studies investigated the relationships across phylogenetic scales (*sensu* Graham et al. 2018), mostly with the aim to decompose biodiversity patterns for higher taxa (Marquet et al. 2004). They typically show that SPRs for different subclades vary and deviate from the pattern observed in the whole large clade (Currie 1991, Buckley et al. 2010, Weiser et al. 2018). Buckley et al. (2010) argue that this finding is inconsistent with the hypotheses explaining richness patterns by productivity. In contrast, Hurlbert and Stegen (2014) argue that weak and varying SPRs of smaller clades are expected if energy availability (productivity) constrains species richness of the whole large clade. Further, Hawkins et al. (2007) show that for many taxa and regions the slope of the STR significantly differs from the slope predicted by the metabolic theory of ecology. However, the patterns for subgroups may differ from those for the whole group due to purely mathematical reasons (Storch and Šizling 2008). Variation of STRs and SPRs across phylogenetic scales is thus likely.

Here, using comprehensive data sets on STRs and SPRs, we simultaneously investigate the effects of multiple factors on their strength and direction. We test several hypotheses, based on above considerations. We expect the strength of both relationships to increase with increasing (1) variance of respective environmental variables, (2) spatial grain and extent and (3) taxon size (total number of species). Additionally, we expect (Fig. 1) that if both temperature and productivity simultaneously affect species richness (Fig. 1b), (4) both STR and SPR should be stronger in regions where temperature correlates positively with productivity, and (5) STRs and SPRs should be similarly strong and approximately equally affected by all the abovementioned factors. In contrast, if only one of these variables directly drives species richness patterns (Fig. 1a), while the other only correlates with it or has an indirect effect (e.g. if temperature affect species richness only by its effect on productivity), then we expect (6) that one of the relationships (either the STR or the SPR) would be more universal, i.e. less variable and less dependent on the examined factors.

**Figure 1.**
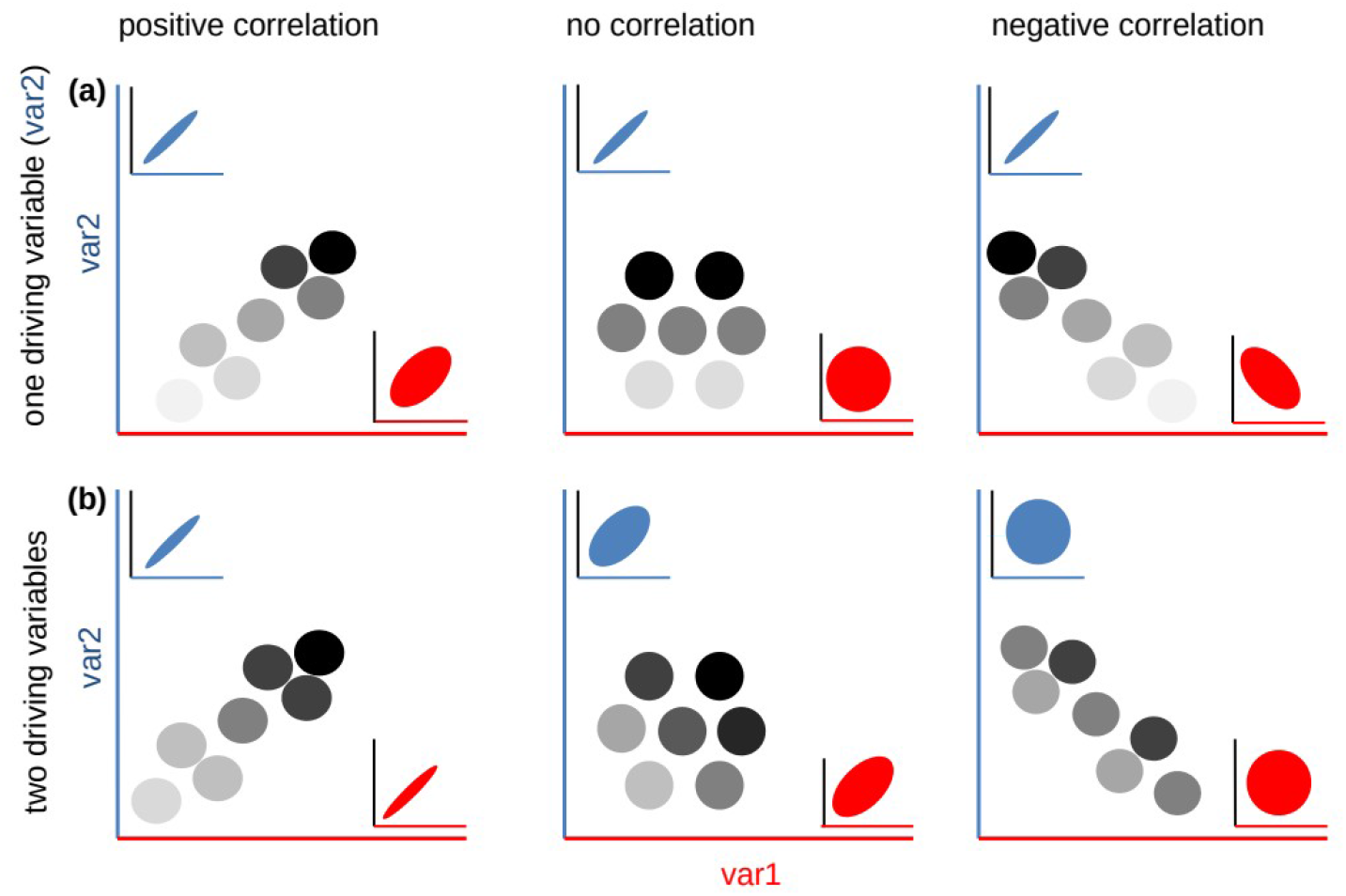
Potential effects of the correlation between two variables (var1 and var2, e.g. temperature and NPP) within a region on the strength of respective relationships of these variables with species richness (e.g. species-temperature and species-productivity relationships). The top row (a) refers to the situation in which only one variable (var2, blue) directly affects species richness, the bottom row (b) to the situation in which both variables independently causally affect species richness. Species richness (circles with the grey scale in the main plot; circle arrangement shows the correlation between var1 and var2) increases from white to black; the ellipses in the insets depict the correlation between respective variable and species richness. In (a) (only var2 has a causal effect), species richness increases only in the vertical direction, and thus the correlation between species richness and var2 is expected to be invariantly tight (and positive), while the strength of the correlation between species richness and var1 depends on the correlation between var1 and var2. In contrast, if both the variables independently affect species richness (b), it increases in the diagonal direction, and the strength of species richness patterns depends (equally for var1 and var2) on the correlation between these variables. The reason is that in this case the variation of the other variable just adds some variation to given relationship.

## Material and methods

### Species data

We used two different types of data for the analysis: (1) a combination of heterogeneous datasets of species richness for different ectothermic taxa spanning different geographical regions (hereafter REGIONS), (2) global species richness data for amphibians, birds and mammals (hereafter CLADES). REGIONS comprise 46 sets of species richness data for a wide range of plant, invertebrate and ectothermic vertebrate taxa used in Hawkins et al. (2007) (Supporting information Table S1). The taxonomic breadth of the datasets ranges from genus (bumblebees) to divisions (vascular plants). The datasets cover all continents (except Antarctica) and their geographical extent ranges from Catalonia (ca. 30,000 km^2^) to global. The datasets also differ in their spatial resolution, most of them using equal-area grid cells (grids from 10 km^2^ to 611,000 km^2^), but in five datasets the units are political regions (Californian plants and butterflies) or nature reserves (Chinese amphibians, reptiles and angiosperms). The datasets were kindly provided by B. Hawkins and more information about the data and their original sources can be found in Hawkins et al. (2007).

In the case of CLADES, we examined the STR and the SPR for every amphibian, avian and mammalian clade (defined by every node) larger than 50 species (to reach sufficient statistical power) within its whole distributional range on the 1° grid. We used the data on the global distribution of mammalian and amphibian species provided by the IUCN Global Species Programme (<http://www.iucnredlist.org>), and avian species provided by BirdLife International (<http://www.datazone.birdlife.org>). These two databases contain distributional information on species ranges in georeferenced polygons for 5,298 terrestrial mammal, 6,493 amphibian, and 10,961 bird species (extinct species were excluded), which we transformed into a 1° (~100.2 km) spatial grid using the Mollweide coordinate system (equal-area projection). We used amphibian phylogeny from <https://doi.org/10.5061/dryad.cc3n6j5> (Jetz and Pyron 2019), avian phylogeny from <http://birdtree.org/> (Jetz et al. 2012) and mammalian phylogeny from <https://doi.org/10.5061/dryad.rc416> (Rosauer et al. 2017).

### Environmental data

For REGIONS we used original temperature data (mean annual temperature) from different sources, assembled by Hawkins et al. (2007). For CLADES we used mean annual temperature data from CHELSA model (Karger et al. 2017, <http://chelsa-climate.org/>, downloaded in April 2017), which we resampled from the original 30 arcsec resolution to the scale of species data. We used net primary productivity (NPP) data averaged from years 2000-2012 compiled from the MODIS NPP layers (MOD17A3, Running et al. 2011, <https://lpdaac.usgs.gov/>) for both REGIONS and CLADES. We resampled NPP data from the original resolution of 1 km to the resolution of each dataset (REGIONS) and to the 1° resolution for CLADES.

### Explanatory variables

We defined several variables describing each dataset (REGIONS) or clade (CLADES), which we then used as the explanatory variables in models explaining variation in the strength of the STR and the SPR. The description of these variables is in Table 1. For the purpose of one of the analyses, we divided the explanatory variables into three categories - phylogenetic scale (*size*), spatial scale (*area* and *grain*) and region characteristics (*mean temperature*, *mean NPP*, *temperature range*, *NPP range* and temperature-NPP correlation – *T_NPP*).

**Table 1.** Explanatory variables for both types of data (REGIONS and CLADES). The abbreviations used in Table 2 are in parentheses in the column VARIABLE. Region characteristics are tinted grey.

| VARIABLE | EXPLANATION | NOTE |
| --- | --- | --- |
| *size* | total number of species in the dataset (REGIONS) or species richness of the clade (CLADES) | unknown for Chinese amphibians, reptiles and angiosperms, Australian tiger beetles, and Iberian/French dung beetles |
| *area* | geographical extent of the dataset in km<sup>2</sup> (REGIONS) or clade range size measured as the number of grid cells occupied by given clade (CLADES) | not defined in New World ants, North American grasshoppers, Chinese amphibians, reptiles and angiosperms (since species richness data did not cover continuously any clearly defined region) |
| *grain* | resolution of the dataset - size of the grid cell for raster data, size of the sampling unit for sample data (New World Ants, North American grasshoppers) and mean area of nature reserves (considering the reserve areas did not vary by more than an order of magnitude) for Chinese amphibians, reptiles and angiosperms in km<sup>2</sup> | defined only for REGIONS (since it is identical for all CLADES); not defined for Californian plants and butterflies because there it represented political units whose area varied by orders of magnitude. |
| *mean temperature (meanT)* | mean annual temperature in °C within the dataset (REGIONS) or within the clade range (CLADES) |  |
| *mean NPP (meanNPP)* | mean NPP within the dataset (REGIONS) or within the clade range (CLADES) |  |
| *temperature range (rangeT)* | range of mean annual temperature within the dataset (REGIONS) or within the clade range (CLADES) (the difference between the lowest and the highest value) |  |
| *NPP range (rangeNPP)* | range of NPP within the dataset (REGIONS) or within the clade range (CLADES) (the difference between the lowest and the highest value) |  |
| *T_NPP* | Spearman's correlation coefficient between temperature and NPP within the dataset (REGIONS) or within the clade range (CLADES) |  |

### Response variables

As a measure of the STR and the SPR strength we used correlation (Spearman’s coefficient) between the number of species and temperature (*S_Temp*) and the number of species and NPP (*S_NPP*), respectively. Spearman’s correlation coefficient reflects both the strength and the direction of the relationships, regardless whether the relationship is linear or curvilinear. Almost all the relationships are monotonic, so the rank-order correlation coefficient well characterizes their strength. However, we performed the analyses also using R-square (coefficient of determination) of the OLS regression model (linear or quadratic, based on AIC comparison) as this measure. It has led to similar results (see the Supporting information). To compare STR strength (*S_Temp*) with SPR strength (*S_NPP*), we calculated mean difference between *S_Temp* and *S_NPP* within each group of data (REGIONS, amphibians, birds, mammals) and performed paired t-tests.

### Statistical analysis

For REGIONS, we used weighted least square (WLS) regression as a basis of all analyses of the strength of STR and SPR, since the frequency distribution of *S_Temp* and *S_NPP* was close to normal. We accounted for different sample size of the datasets by using weighting by square root of sample size. First, we performed individual regression models for each single explanatory variable alone. We tested for the curvilinearity in these univariate models by comparing the AIC of models containing the explanatory variable in (1) linear term and in (2) both linear and quadratic term. If the AIC difference was lower than 2, we considered the relationship linear, otherwise we used the model with lower AIC (Burnham and Anderson 2002). Second, we explored how much variation in *S_Temp* and *S_NPP* was explained by the full model using all explanatory variables (in linear term) together. We are aware of the fact that these full models are not entirely statistically accurate due to collinearity in data. Still, since we only wanted to estimate the total variation explained by all variables (i.e. in this case we did not explore model parameters comprising the effects of individual variables), since we present the adjusted R^2^ values (which adjust for the number of predictors), and since we deal with the collinearity in further analysis, we consider this approach appropriate. Third, we used variation partitioning analysis to distinguish independent effects of the explanatory variables. We used adjusted R^2^ to assess the variation explained by the explanatory variables and their combinations, because it is the only unbiased method (Peres-Neto et al. 2006). To keep the results clear, we did not use all the individual explanatory variables in the partitioning. Instead, we first performed variation partitioning for three groups of variables - (i) phylogenetic scale (variable *size*), (ii) spatial scale (variables *grain* and *area*) and (iii) region characteristics (the three region characteristics which had the strongest effects in the univariate models). Then, we further decomposed the effects of these three region characteristics. Each variable entered the variation partitioning analysis in linear or quadratic form depending on the results of the univariate models. Variables *size*, *area* and *grain* were log-transformed in order to linearise the observed relationships.

For CLADES, to account for the phylogenetic correlation and nestedness among our data points (clades), we used phylogenetic generalized least squares (PGLS, Freckleton et al. 2002) as a basis of all analyses. Specifically, we first estimated phylogenetic dependence among the clades by searching for the maximum likelihood value of the parameter λ, which varies from 0 (phylogenetic independence) to 1 (phylogenetic signal consistent with the Brownian evolutionary model), and then we fitted a GLS model while controlling for the phylogenetic dependence, using package ‘caper’ (https://CRAN.R-project.org/package=caper, Orme et al. 2013, Machac et al. 2018). This way we performed all the analyses described above - univariate models, full model and the variation partitioning analysis. We performed all the analyses for 100 randomly selected phylogenetic trees and then averaged the results from these 100 models. Log-transformation was used for variables *size* (for all three vertebrate groups) and *area* (for amphibians) in order to linearise the observed relationships.

PGLS accounts for the dependence among clades given by their phylogenetic distance, but not necessarily for their nestedness, causing that patterns observed within a clade may be affected by patterns concerning the subclades and vice versa (Graham et al. 2018). There is no simple solution of this problem. One possibility would be to use only a set of non-nested clades (Machac et al. 2018). Nevertheless, we preferred not to use this approach, as the selection of exclusively nonnested clades would be highly non-random, would lead to small sample sizes, and would require repeated analyses on different non-nested clades sets, which would complicate the results interpretation. Using all possible clades circumvents the problem of clade selection, and also involves a range of clades, from lower to higher taxa, providing more complex results. Results from other studies (Machac et al. 2018) indicate that the results for nested and non-nested clades are similar, and although nestedness may bias statistical significance of the tests, it does not affect the patterns themselves. All the analyses were performed in R (R Development Core Team 2017).

## Results

### Strength of STRs and SPRs

NPP is on average a stronger correlate of species richness than temperature for all tested data (Fig. 2a and 2b). STRs are negative (and concurrently strong) for many datasets and vertebrate clades, while SPRs are in the majority of cases positive (e.g. for REGIONS, STRs are negative for 16 datasets, while SPRs for 5 datasets and they are weak for most of these negative values). Since negative STR is not an evidence for temperature as a driving factor of species richness patterns, NPP can be considered a much better predictor of species richness than temperature. Specifically, *S_NPP* is on average by 0.21 higher than *S_Temp* for REGIONS (t_45_=-2.48, p=0.02), by 0.30 higher for amphibians (t_360_=-14.49, p<10^-16^), by 0.19 higher for birds (t_501_=-14.53, p<10^-16^) and by 0.20 higher for mammals (t_248_=-9.24, p<10^-16^) (paired t-tests). The highest proportion of negative STRs and related highest mean difference between *S_Temp* and *S_NPP* found for amphibians likely results from their dependency on water availability. Still, for many datasets and clades, temperature is a better predictor of species richness than NPP, although sometimes both variables and sometimes neither variable predicts species richness well (Fig. 2c).

**Figure 2.**
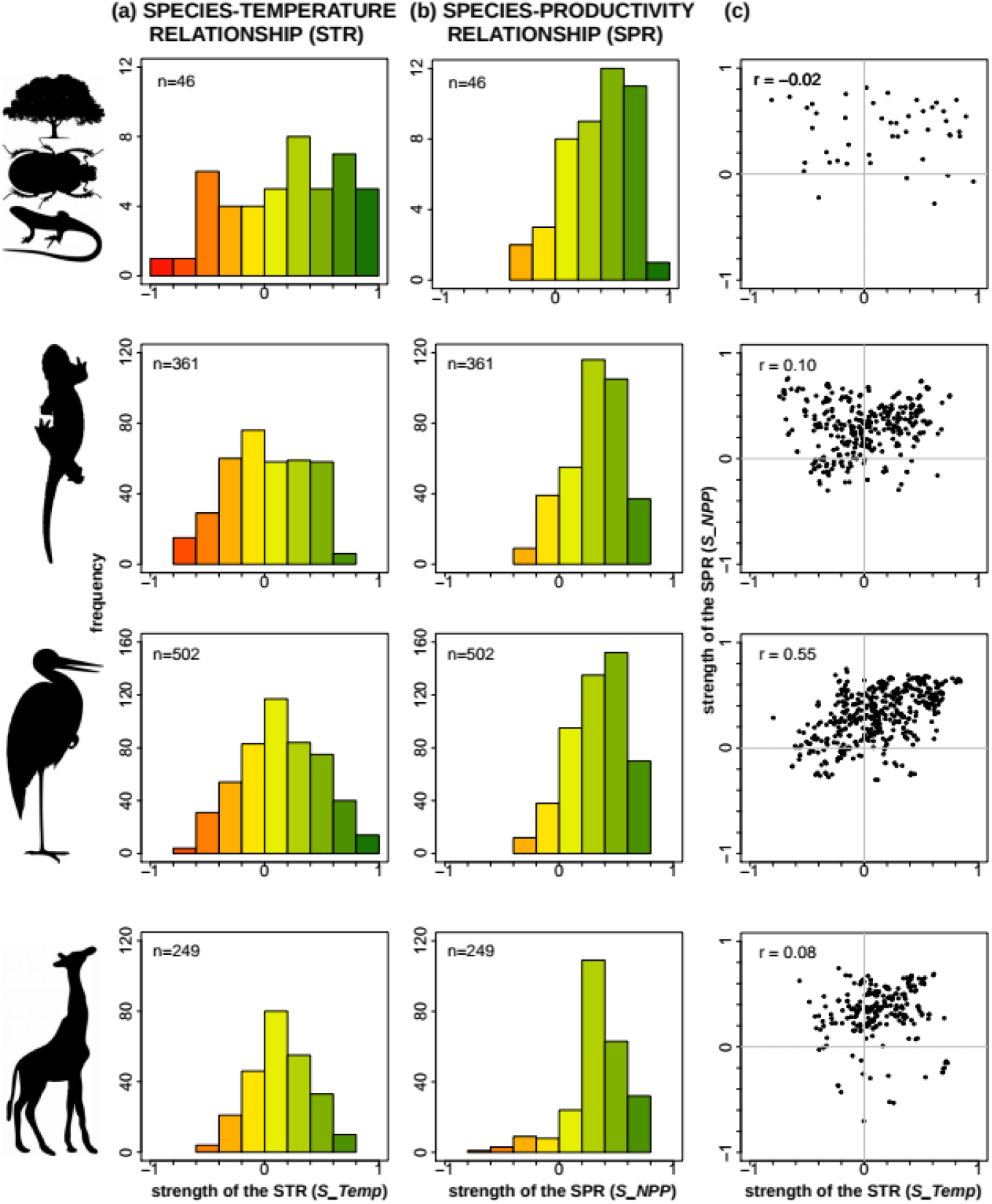
Histograms showing the distribution of the frequencies of (a) the STR strength and (b) the SPR strength for each data set (the first row refers to REGIONS, the following three rows to CLADES). The colours are scaled from yellow (weak relationship) to red (negative relationship) and to green (positive relationship) and correspond to the colour scale in Fig. 3. Sample size (n) is presented inside each plot. (c) The relationship between the strength of the STR (*S_Temp*) and the strength of the SPR (*S_NPP*) for each data set (rows). The Pearson’s correlation coefficient (r) is presented inside each plot.

### Factors affecting strength of STRs and SPRs in REGIONS

In REGIONS, we are able to explain 55% of variation in the strength of STRs (*S_Temp*) and 13% of variation in the strength of SPRs (*S_NPP*) by our explanatory variables (adjusted R^2^ of the WLS full model), indicating that the SPR is less dependent on the examined factors. *S_Temp* strongly depends on *mean temperature*, on *T_NPP* and on *temperature range* (Table 2a). Specifically, STRs are generally stronger in colder regions, in regions where temperature and NPP are positively correlated and in regions with wider range of temperature (Table 2a, Fig. 3, Supporting information Fig. S1). On the other hand, the strength of the SPR is not as well predictable by our explanatory variables (Table 2a). *S_NPP* depends on *mean temperature*, but in the opposite direction than *S_Temp*, and there is a significant effect of *T_NPP* (Table 2a). SPRs are generally stronger in warmer regions and in regions where temperature and NPP are correlated (both positively and, curiously, negatively) (Table 2a, Fig. 3, Supporting information Fig. S1).

**Figure 3.**
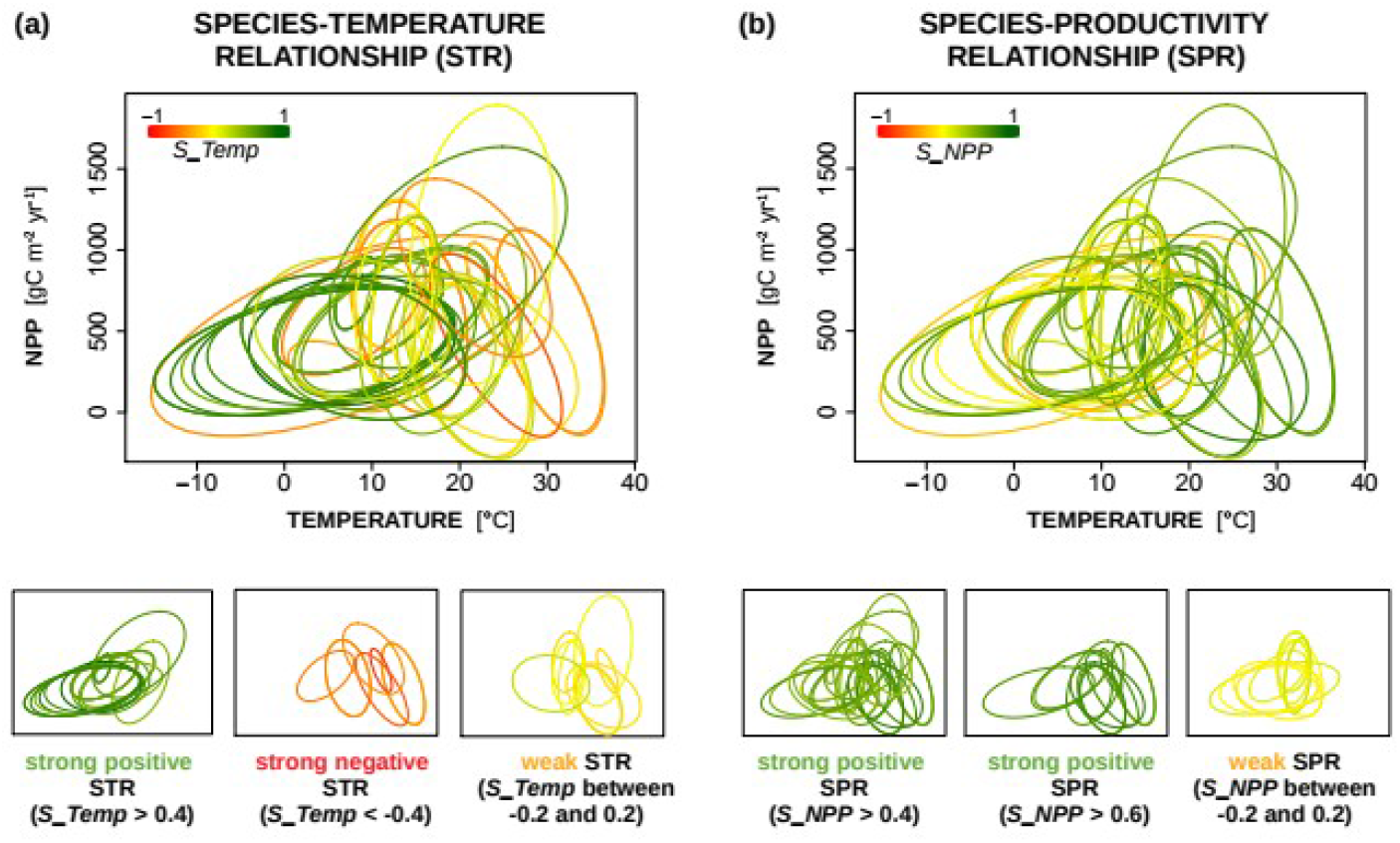
Plots showing the strength of (a) the STR and (b) the SPR of REGIONS in relation to region characteristics. Mean values and ranges of temperature and NPP are reflected by the position of the ellipse within the plot, and the temperature-NPP correlation is reflected by the diagonal elongation of the ellipse. Each ellipse represents one dataset and it encompasses 80% of data points. Colours of the ellipses correspond to the strength of the STR (*S_Temp*) and the SPR (*S_NPP*), respectively, according to the colour bar. Additionally, we split the datasets so that only the strong positive, strong negative and weak relationships are presented (bottom). Note that there are no strong negative SPRs. (a) STRs are strong and positive in colder regions, regions with wider temperature range and regions with positive temperature-NPP relationship. On the other hand, STRs are strong and negative in warmer (tropical) regions, in regions with narrower temperature range and those revealing negative temperature-NPP relationships. They are generally weak for regions where temperature and NPP are not correlated. (b) SPRs do not show as clear patterns as STRs, although the effect of the mean temperature is still visible – SPRs are generally stronger in warmer regions and they are mostly weak in regions where temperature and NPP are not correlated.

**Table 2.**
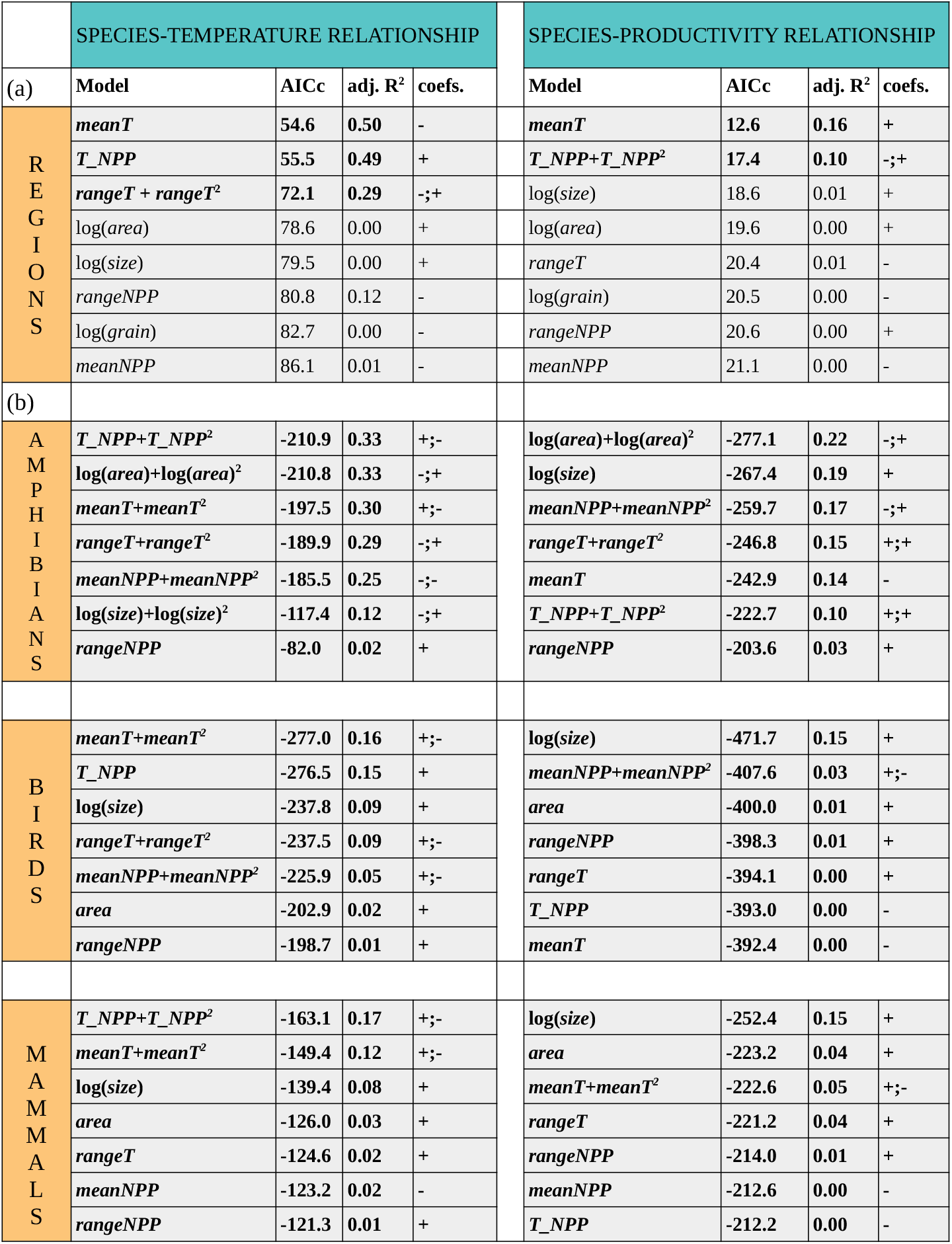
Results of the univariate models explaining the variation in the strength of the STR (left column) and in the strength of the SPR (right column) for (a) datasets of various ectothermic taxa – REGIONS, and for (b) vertebrate clades – CLADES. The effects significant on α=0.05 are in bold, for the explanation of the variables see Table 1. AIC corrected for small sample size (AICc), adjusted *R^2^* (adj. *R^2^*) and coefficients (coefs.) are displayed.

Interestingly, the strength of STRs and SPRs for REGIONS depends neither on taxon size nor on spatial scale (in terms of both *grain* and *area*; Table 2a). The absence of a univariate effect of the geographical extent (*area*) suggests that environmental gradients (namely *temperature range*) are more important determinants of the strength of the STR than geographical extent per se. Indeed, the variation partitioning indicates that when we compare different taxa in different (arbitrarily chosen) regions of the world, the strength of the STR and the SPR depends almost entirely on environmental characteristics of the selected region (Fig. 4), although *mean temperature* and *T_NPP* are correlated (Fig. 3a) and we cannot completely separate their effects. In the case of STRs (*S_Temp*), the largest portion of variation is explained by the joined effects of *mean temperature* and *T_NPP*, but there is also a separate effect of *T_NPP* (unlike *mean temperature*, Fig. 4a). In contrast, in the case of SPRs (*S_NPP*), the separate effect of *mean temperature* is larger than the separate effect of *T_NPP* (Fig. 4b). Thus, the strength of SPRs depends mostly on mean temperature within the region (SPRs being stronger in warm, low latitude regions) but the strength of STRs depends on how temperature correlates with NPP within the region (in accord with Fig. 1a; compare predictions in Fig. 1a with data in Fig. 3a), suggesting that the NPP is a more proximate driver of species richness than temperature.

**Figure 4.**
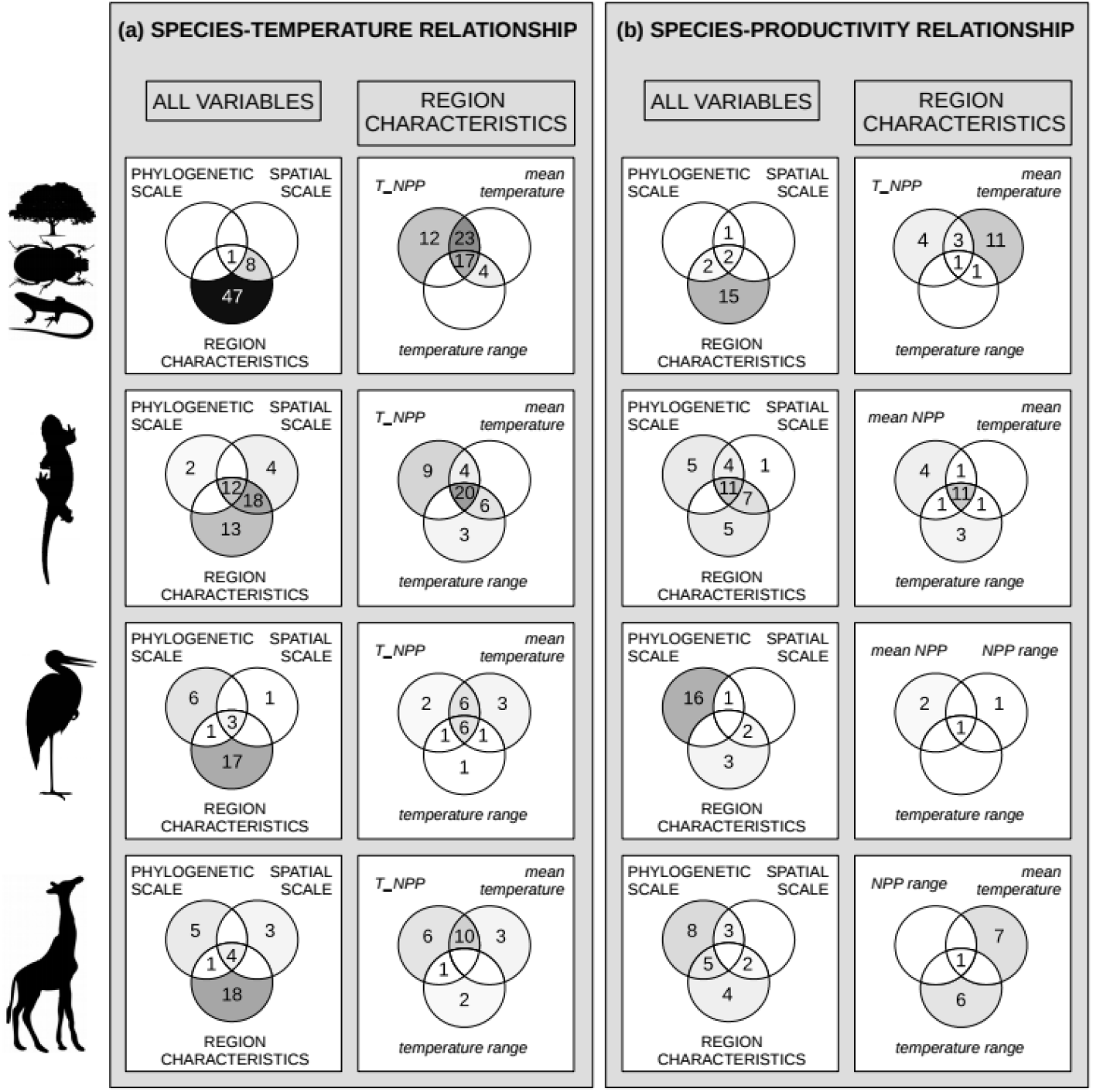
Venn diagrams describing variation partitioning for (a) the strength of the STR (*S_Temp*) and (b) the strength of the SPR (*S_NPP*) for different data (rows). Within each panel, in the left column, explanatory variables are divided into 3 categories - phylogenetic scale (variable *size*), spatial scale (variables *area* and *grain [grain* only for REGIONS]) and region characteristics (three region characteristics with the strongest univariate effect on *S_Temp* or *S_NPP*, respectively, see Table 2 and Table 1). In the right column, the variation partitioning is further performed for these three region characteristics. The numbers represents adjusted *R^2^* (in %), zero and negative values are not displayed. Every section is coloured according to its adjusted *R^2^* value on the white-black scale (white corresponds to 0, black corresponds to 50). *Note: T_NPP* abbreviates the temperature-NPP correlation.

### Factors affecting strength of STRs and SPRs in CLADES

For CLADES, the results are different in several respects. First, our variables explain lower amount of variation in the strength of STRs than in REGIONS, especially for avian and mammalian clades. The full models explain 44% of the variation in STRs strength and 32% of the variation in SPRs strength for amphibians, 21% of the variation in STRs strength and 20% of the variation in SPRs strength for birds, and 17% of the variation in STRs strength and 23% of the variation in SPRs strength for mammals (adjusted R^2^ of the PGLS full models). Second, in contrast to REGIONS, there is a significant effect of the clade size on the strength of the relationships (Table 2b), indicating that the expected effect of the taxon size is important only when comparing different subclades within one taxonomical group. Moreover, taxon size is the only important factor explaining the strength of SPRs in the univariate models in avian and mammalian clades (Table 2b). However, even the univariate effect of taxon size is not very strong (Table 2b), as STRs and SPRs are more idiosyncratic for smaller clades, while for large clades they are strong in vast majority of cases (Supporting information Fig. S2 and S3). Third, only amphibians reveal similarly strong univariate effects of region characteristics on the STR strength as in the case of REGIONS. The endothermic taxa (birds and mammals) reveal weaker effects of all the explanatory variables, although *T_NPP* and *mean temperature* remain the strongest determinants of the STR strength (Table 2b, Supporting information Fig. S2).

The effect of phylogenetic scale (clade size) is shown also by the variation partitioning analysis, where partial effects of clade size are ubiquitous (Fig. 4, note that ‘phylogenetic scale’ represents only one variable, i.e. *size*, while ‘region characteristics’ represents three variables). Variation partitioning analysis also reveals that partial effects of the region characteristics are still important for the STR while they are weak for the SPR (Fig. 4).

## Discussion

We provide the first analysis that compares the effects of multiple taxon- and region-related variables on the strength of the STR and the SPR. We investigated this using two different types of data, representing two common types of data structure in macroecological research: (1) various taxa sampled using different sampling techniques within different regions of the world (Hawkins et al. 2007) and (2) grid-cell data of amphibian, avian and mammalian clades within their global ranges. Two major findings have emerged. First, spatial scale and taxon size are relatively unimportant determinants of the strength of both STR and SPR in comparison with the characteristics of the region within which species richness data are sampled. Second, species-temperature relationships are context-dependent, more variable and weaker than species-productivity relationships, indicating that the role of temperature in biodiversity patterns is of secondary importance in comparison with the role of productivity, which is quite universal determinant of species richness. Below we discuss these findings and their implications in detail.

### The effects of taxon size and spatial scale on the STR and the SPR

There was a striking difference in the drivers of variation in the explored species richness patterns between the two types of data. Taxon size affected the strength of STRs and SPRs only when comparing subclades within vertebrate classes, i.e. when exploring these relationships across different phylogenetic scales (sensu Graham et al. 2018). Weiser et al. (2018) in their study analysing latitudinal gradient of species richness (LGSR) for different nested plant clades concluded that this non-universality and variability of LGSR suggests that climate or energy fail to explain LGSR and that this result instead supports the ‘tropical conservatism’ hypothesis. However, in accord with Hurlbert and Stegen (2014), we argue that this inconsistency in smaller taxa does not exclude temperature or productivity as important factors shaping species richness patterns. There are several possible reasons why smaller taxa may exhibit weak and variable relationships: (1) simple statistical reasons (sampling effects), (2) species richness of small clades may be affected by a specific environmental factor related to specifities of their biology, or interaction with other taxa, (3) smaller taxa may be young and their richness patterns may be thus driven by their recent history of spreading from an area of origin (Gehrke and Linder 2011).

In this respect, it is interesting that we did not find any univariate effects of clade size when comparing different taxa within different regions (i.e. the meta-analytic approach for REGIONS; as in e.g. Mittelbach et al. 2001, Hawkins et al. 2007, Field et al. 2009). One explanation could be that 46 datasets do not represent sufficient statistical power for revealing such an effect. However, we were able to show other strong effects using the same data, so the statistical power does not seem to be an issue. Therefore, a more plausible explanation is that when comparing different taxa in different regions, it only matters which region is selected from the global map of temperature-productivity conditions, relatively independently of the taxon studied. The regions are delimited arbitrarily and almost never represent real ranges of the whole taxa, so that the taxon size becomes unimportant in comparison with the environment (mean temperature, temperature-productivity relationship or the range of temperature within the region). This explains also the absence of univariate effects of spatial scale (both grain and extent).

### Comparison between the species-temperature and species-productivity relationship

We have shown that STRs are weaker and more variable than the SPRs, and that STR variation is attributable to the characteristics of the region, most importantly to the correlation between temperature and productivity (in accord with Fig. 1a). STRs are also stronger if the region encompasses wider temperature range (especially in ectotherms), which is probably a simple statistical effect – pronounced STR requires sufficient temperature variation. The observation that STR is often not particularly strong and varies widely, sometimes being even negative, may not seem particularly surprising, given that high temperature is associated with high species richness only if water is not limited – hot arid regions host low number of species due to the scarcity of resources. Anyway, it means that temperature cannot be considered the main factor affecting species richness on land. This is supported by our finding that the strength of the STR is mostly affected by the temperature-productivity correlation (which is quite evenly distributed from strong negative to strong positive in our data, Supporting information Fig. S1, S2 and S3). Temperature may thus affect species richness only indirectly, via its positive effect on productivity (Šímová and Storch 2017) and/or the length of the vegetation season, which affects temporal availability of resources. In both these cases, resource availability would be the proximate driver of species richness patterns.

The effect of temperature on the length of the growing season may also explain our finding that, in contrast to SPRs which are generally stronger in warmer regions (within the tropics), STRs are stronger in colder regions, i.e. within temperate zone with pronounced seasonality. This observation is consistent with earlier studies showing that in higher latitudes, species richness is best predicted by temperature or potential evapotranspiration (PET, driven mostly by temperature), while in lower latitudes it is best explained by productivity or water availability (Hawkins et al. 2003a). Similarly, avian species diversity was shown to be best predicted by temperature or PET within Palearctic and Nearctic regions, and by productivity or rainfall in the Afrotropics, Neotropics and Australia (Hawkins et al. 2003b). On the other hand, we cannot exclude the possibility that the effect of temperature on species richness goes beyond its simple effect on resource abundance and/or its temporal availability. If temperature affected only productivity or the length of the growing season, STRs should be in all cases weaker than SPRs, since mean annual productivity should reflect the length of the season as well. Although the general patterns agree with this expectations, there are exceptions (Fig. 2c), STRs being sometimes stronger than SPRs. This may be partly attributable to the fact that productivity is more difficult to measure or estimate than temperature (Šímová and Storch 2017). However, other effects of temperature may still play a role. The finding that temperature has a stronger effect in temperate regions may be interpreted as an effect of limited pool of available species that can tolerate lower temperatures (Šímová et al. 2011), in accord with the niche conservatism hypothesis (Wiens and Donoghue 2004). Additionally, low temperature may limit diversification rates in ectotherms (Allen et al. 2007, Worm and Tittensor 2018).

Regardless of these considerations, productivity appears as more important factor driving species richness patterns. Although SPRs were also a bit stronger in regions where productivity and temperature positively correlated, the effect was quite weak and mostly shared with the effect of mean temperature (Fig. 4b). Moreover, the regions in which temperature and productivity were negatively correlated (typically in the tropics; see Fig. 3) revealed negative STRs but positive SPRs. The strength of species-productivity relationships was even less dependent on all the explored variables in endotherms. While SPR variation in amphibian clades reveals similar causes as the variation in the datasets of various ectothermic taxa (REGIONS) (with the exception of differences caused by the different nature of the data discussed above), the SPRs for avian and mammalian clades are largely independent of our explanatory variables, with the exception of the taxon size discussed above. Given that SPRs are generally strong (Fig. 2) and independent of most of the examined variables in endotherms, productivity indeed appears as the key factor shaping species richness patterns for these taxa. Although temperature may be as good predictor of species richness as NPP for some clades and datasets (Fig. 2c), the finding that *in general* SPRs are positive, strong, and largely independent of most of the explored variables (compared to STRs, Fig. 2) indicates that productivity is more important species richness driver than temperature - and this is especially pronounced in endotherms.

### Conclusion

We provide the first comprehensive test of how different variables comprising spatial scale and the characteristics of examined taxa and regions affect the strength of species-temperature and species-productivity relationships. Both relationships are stronger for larger clades, indicating that phylogenetic scale matters (Weiser et al. 2018), but only when comparing different subclades within one large taxon. When comparing different taxa in different regions (meta-analytical approach), only the role of region characteristics (mean temperature and temperature-productivity correlation within the region) has been revealed. We provide new evidence for the previously documented patterns that temperature plays a more important role in colder regions, while productivity is more important in warmer regions (Hawkins et al. 2003a, 2003b). Additionally, species-temperature relationships are weak if temperature does not correlate with productivity. The relationships between species richness and productivity are less variable and at the same time more independent of all the studied factors, indicating that the species-productivity relationship is more universal and context-independent than the species-temperature relationship. This may stem from the universal productivity effects on the limits of species richness (sensu Hurlbert and Stegen 2014), i.e. species richness carrying capacity (sensu Storch and Okie 2019). We suggest that every study aiming to test a mechanism hypothesised to form spatial patterns in species richness should pay careful attention to the effects potentially causing the variation of the strength of species richness patterns, especially when data only for arbitrarily chosen regions and/or taxa are used.

## Data availability statement

Most of the data used in the study (with the exception specified below) are freely available from the cited and publicly accessible databases. 46 datasets containing data on species richness and mean annual temperature for different taxa and regions (referred as REGIONS in the text) were assembled by Hawkins et al. (2007) and kindly provided by Bradford Hawkins for the purpose of the study. We provide the list of the datasets with the values of response and explanatory variables used in our analysis in the Supporting information.

