## Supporting information for "Universality in biodiversity patterns: Variation in species-temperature and species-productivity relationships reveals a prominent role of productivity in diversity gradients"

**Table S1.** List of the datasets (our basic units, referred as REGIONS) used in the analysis with the number of samples and the values of response and explanatory variables. *S_Temp* - strength of the STR (Spearman's correlation coefficient between the number of species and temperature), *S_NPP* - strength of the SPR (Spearman's correlation coefficient between the number of species and NPP), *S_Temp.R2* (used only in the supporting analysis) - strength of the STR (coefficient of determination of the OLS regression model explaining number of species by temperature), *S_NPP.R2* (used only in the supporting analysis) - strength of the SPR (coefficient of determination of the OLS regression model explaining number of species by NPP), *size* - number of species in the dataset, *area* - spatial extent of the dataset, *meanT* - mean temperature within the dataset, *meanNPP* - mean NPP within the dataset, *rangeT* - temperature range within the dataset, *rangeNPP* - NPP range within the dataset, *T_NPP* - Spearman's correlation coefficient between temperature and NPP within the dataset, *grain* - spatial resolution of the dataset (detailed description of the variables is in Table 1).

| **DATASET** | **NUMBER OF SAMPLES** | **RESPONSE VARIABLES** | | | | **EXPLANATORY VARIABLES** | | | | | | | |
|---|---|---|---|---|---|---|---|---|---|---|---|---|---|
| | | ***S_Temp*** | ***S_NPP*** | ***S_Temp.R2*** | ***S_NPP.R2*** | ***size*** | ***area*** | ***meanT*** | ***meanNPP*** | ***rangeT*** | ***rangeNPP*** | ***T_NPP*** | ***grain*** |
| Afrotropical amphibians | 1813 | -0.455 | 0.661 | -0.415 | 0.522 | 615 | 19988325 | 30.30 | 500.65 | 20.31 | 1657.45 | -0.578 | 105 |
| Afrotropical snakes | 1948 | -0.501 | 0.626 | -0.463 | 0.442 | 406 | 21476700 | 30.64 | 493.35 | 20.31 | 1657.45 | -0.596 | 105 |
| Australian amphibians | 604 | 0.079 | 0.670 | 0.129 | 0.403 | 223 | 7308400 | 21.08 | 258.94 | 21.25 | 1822.04 | -0.344 | 110 |
| Australian butterflies | 625 | -0.160 | 0.755 | -0.157 | 0.649 | 393 | 7562500 | 21.47 | 258.87 | 20.60 | 1822.04 | -0.328 | 110 |
| Australian tiger beetles | 53 | 0.295 | 0.354 | 0.114 | 0.047 | NA | 7227500 | 21.17 | 290.31 | 15.70 | 1553.23 | -0.410 | 350 |
| Brasilian cerrado amphibians | 178 | -0.811 | 0.699 | -0.661 | 0.564 | 131 | 2189400 | 23.90 | 709.93 | 8.67 | 1028.91 | -0.687 | 111 |
| Brasilian cerrado reptiles | 181 | -0.136 | 0.276 | -0.147 | 0.293 | 706 | 2189400 | 23.90 | 712.37 | 8.67 | 1028.91 | -0.682 | 111 |
| Californian butterflies | 90 | -0.528 | 0.026 | -0.309 | 0.014 | 217 | 423970 | 14.40 | 587.67 | 18.94 | 1486.58 | -0.317 | NA |
| Californian plants | 90 | -0.418 | 0.574 | -0.459 | 0.515 | 5902 | 423970 | 14.40 | 587.67 | 18.94 | 1486.58 | -0.317 | NA |
| Catalonian Orthoptera | 301 | -0.303 | 0.110 | -0.102 | 0.012 | 161 | 30100 | 12.32 | 845.11 | 12.99 | 1467.81 | -0.024 | 10 |
| Catalonian plants | 184 | -0.150 | 0.097 | -0.043 | 0.032 | 3087 | 28500 | 12.35 | 850.65 | 11.53 | 1461.50 | -0.048 | 10 |
| Coloradan/Nevadan ants | 56 | -0.331 | 0.206 | -0.052 | 0.161 | 226 | 540792 | 6.57 | 221.18 | 17.70 | 469.81 | -0.488 | 110 |
| European amphibians | 386 | 0.517 | 0.593 | 0.619 | 0.405 | 49 | 4670600 | 8.52 | 603.38 | 26.51 | 1775.64 | 0.422 | 110 |
| European pteridophytes | 2257 | -0.518 | 0.107 | -0.287 | 0.007 | 158 | 5642500 | 8.09 | 622.06 | 22.76 | 1731.43 | 0.481 | 50 |
| European reptiles | 386 | 0.754 | 0.364 | 0.610 | 0.229 | 71 | 4670600 | 8.52 | 603.38 | 26.51 | 1775.64 | 0.422 | 110 |
| European trees | 386 | 0.559 | 0.419 | 0.544 | 0.295 | 187 | 4670600 | 8.51 | 603.38 | 26.51 | 1775.64 | 0.422 | 110 |
| Great Britain plants (exotics) | 2250 | 0.834 | 0.400 | 0.659 | 0.366 | 1592 | 225000 | 8.19 | 734.84 | 7.58 | 1137.41 | 0.477 | 10 |
| Great Britain plants (natives) | 2250 | 0.720 | 0.500 | 0.520 | 0.345 | 1462 | 225000 | 8.19 | 734.84 | 7.58 | 1137.41 | 0.477 | 10 |
| Global bumble bees | 143 | -0.399 | -0.222 | -0.454 | -0.068 | 239 | 87373000 | 6.85 | 473.46 | 45.00 | 1740.29 | 0.565 | 782 |

| | | | | | | | | | | | | | |
|---|---|---|---|---|---|---|---|---|---|---|---|---|---|
| Chinese amphibians | 54 | 0.634 | 0.675 | 0.403 | 0.556 | NA | NA | 12.17 | 558.92 | 26.70 | 1280.56 | 0.668 | 13 |
| Chinese angiosperms | 51 | 0.367 | 0.399 | 0.411 | 0.190 | NA | NA | 12.11 | 539.00 | 24.70 | 1280.56 | 0.587 | 13 |
| Chinese reptiles | 54 | 0.598 | 0.630 | 0.380 | 0.313 | NA | NA | 12.46 | 565.41 | 24.70 | 1280.56 | 0.662 | 12 |
| Iberian amphibians | 257 | 0.232 | 0.484 | 0.010 | 0.156 | 30 | 642500 | 13.77 | 631.45 | 13.13 | 1620.72 | 0.286 | 50 |
| Iberian seed plants | 257 | -0.233 | 0.125 | -0.124 | 0.076 | 2687 | 642500 | 13.77 | 631.45 | 13.13 | 1620.72 | 0.286 | 50 |
| Iberian reptiles | 257 | 0.043 | 0.182 | -0.002 | 0.008 | 50 | 642500 | 13.77 | 631.45 | 13.13 | 1620.72 | 0.286 | 50 |
| Indian tiger beetles | 57 | 0.020 | 0.816 | 0.000 | 0.672 | 161 | 4310625 | 22.22 | 329.74 | 30.50 | 1386.10 | -0.310 | 275 |
| Kenyan woody plants | 157 | -0.652 | 0.729 | -0.384 | 0.465 | 1417 | 492352 | 24.14 | 413.26 | 18.15 | 1528.98 | -0.707 | 56 |
| Mexican hawk moths | 173 | 0.461 | 0.699 | 0.293 | 0.563 | 820 | 1964704 | 19.33 | 517.86 | 20.60 | 1749.77 | 0.399 | 110 |
| North American amphibians | 1444 | 0.892 | 0.543 | 0.793 | 0.324 | 221 | 17472400 | 2.91 | 373.09 | 39.20 | 1368.57 | 0.434 | 110 |
| North American blister beetles | 190 | 0.615 | -0.279 | 0.362 | -0.128 | 173 | 12739416 | 10.63 | 476.90 | 25.50 | 1360.06 | 0.130 | 278 |
| North American butterflies (summer) | 1443 | 0.837 | 0.357 | 0.627 | 0.191 | 578 | 17460300 | 2.16 | 372.91 | 44.43 | 1368.57 | 0.453 | 110 |
| North American butterflies (winter) | 1443 | 0.748 | 0.371 | 0.533 | 0.197 | 535 | 17460300 | 2.16 | 372.91 | 44.43 | 1368.57 | 0.453 | 110 |
| North American grasshoppers | 52 | 0.514 | 0.139 | 0.351 | 0.033 | 305 | NA | 6.92 | 429.95 | 37.00 | 930.26 | 0.405 | 63 |
| North American reptiles | 1444 | 0.957 | -0.070 | 0.900 | -0.108 | 224 | 17472400 | 2.91 | 373.09 | 39.20 | 1368.57 | 0.434 | 110 |
| North American tiger beetles | 192 | 0.734 | -0.012 | 0.662 | 0.040 | 93 | 14520000 | 5.86 | 418.89 | 36.60 | 950.65 | 0.147 | 275 |
| North American trees | 1444 | 0.808 | 0.699 | 0.671 | 0.502 | 676 | 17472400 | 2.91 | 373.09 | 39.20 | 1368.57 | 0.434 | 110 |
| New World ants | 66 | 0.696 | 0.592 | 0.581 | 0.415 | 911 | NA | 18.55 | 953.72 | 22.45 | 1640.73 | 0.524 | 1 |
| Southern African woody plants | 130 | 0.207 | 0.766 | 0.088 | 0.558 | 1372 | 2600000 | 19.31 | 378.42 | 11.81 | 1027.31 | 0.267 | 141 |
| Southern African reptiles | 281 | 0.247 | 0.358 | 0.014 | 0.070 | 664 | 3400100 | 19.10 | 410.94 | 14.58 | 1182.38 | 0.200 | 110 |
| South American tiger beetles | 132 | 0.154 | 0.525 | 0.155 | 0.330 | 264 | 5104688 | 22.42 | 1031.01 | 20.58 | 2195.84 | 0.008 | 137.5 |
| Southeast Asian hawk moths | 430 | -0.454 | 0.435 | -0.219 | 0.270 | 382 | 1300750 | 21.73 | 890.29 | 33.40 | 1708.74 | -0.682 | 55 |
| Iberian pteridophytes | 240 | 0.389 | 0.546 | 0.128 | 0.244 | 114 | 600000 | 13.09 | 626.42 | 13.00 | 1620.72 | 0.430 | 50 |
| Iberian/French dung beetles | 155 | 0.053 | 0.103 | 0.062 | 0.007 | NA | 500000 | 12.51 | 682.98 | 10.16 | 1420.86 | 0.109 | 50 |
| Western Palearctic butterflies | 162 | -0.165 | 0.531 | -0.559 | 0.290 | 413 | 7840800 | 9.50 | 517.70 | 30.05 | 1515.92 | 0.008 | 220 |
| Western Palearctic Eupelmidae | 469 | 0.281 | 0.481 | 0.590 | 0.186 | 150 | 25467645 | 11.69 | 401.16 | 37.28 | 1366.89 | -0.182 | 278 |
| Western Palearctic dung beetles | 457 | 0.373 | -0.038 | 0.558 | -0.078 | 534 | 24900451 | 11.75 | 402.08 | 37.25 | 1366.89 | -0.185 | 278 |

**Figure S1.** The relationships between (a) the strength of the STR (*S_Temp*) and (b) the strength of the SPR (*S_NPP*) and those explanatory variables that significantly affect the STR strength and the SPR strength for REGIONS. Grey lines represents univariate models in Table 2a, detailed description of the variables is in Table 1.

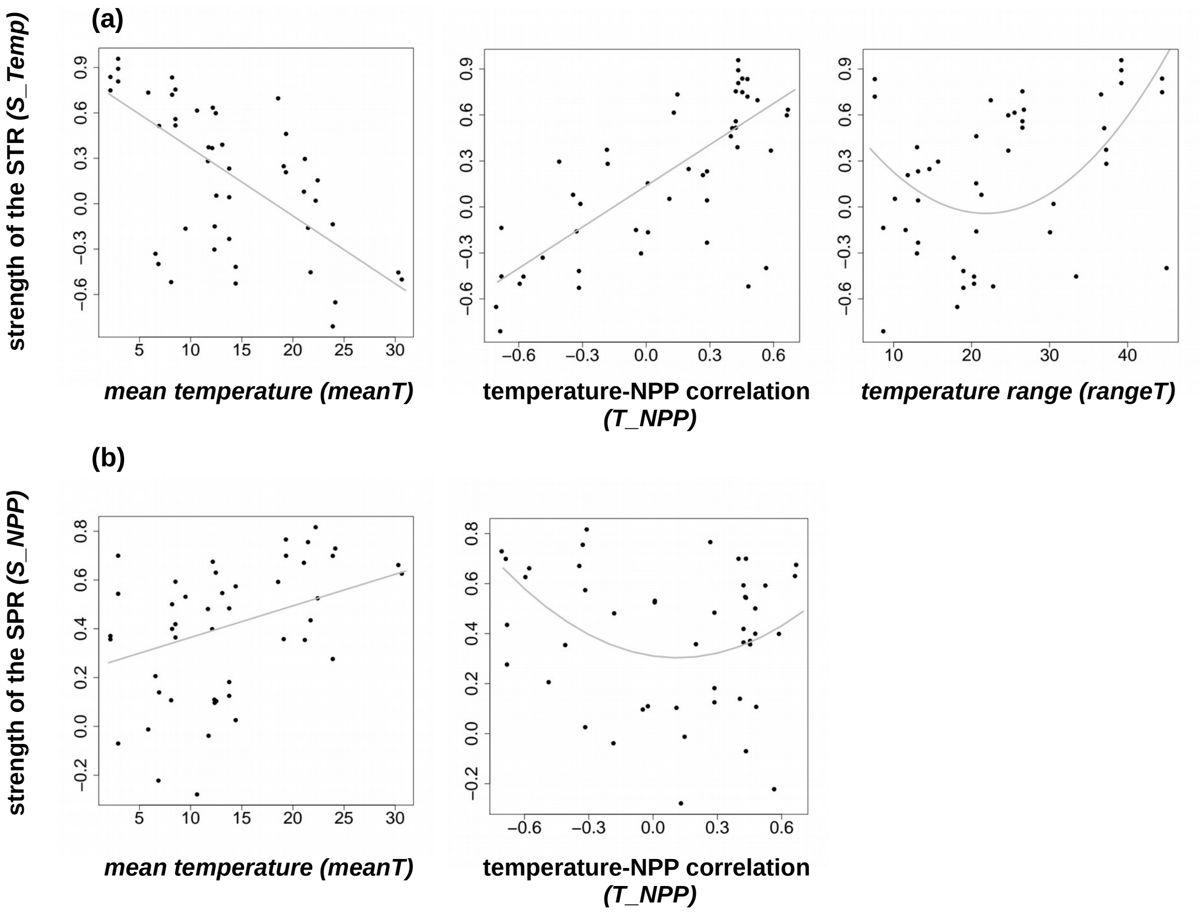

**Figure S2.** The relationships between the STR strength (*S_Temp*) and each explanatory variable (rows) for CLADES (all the PGLS univariate models are significant, see Table 2b). Grey lines represents univariate models in Table 2b, detailed description of the variables is in Table 1.

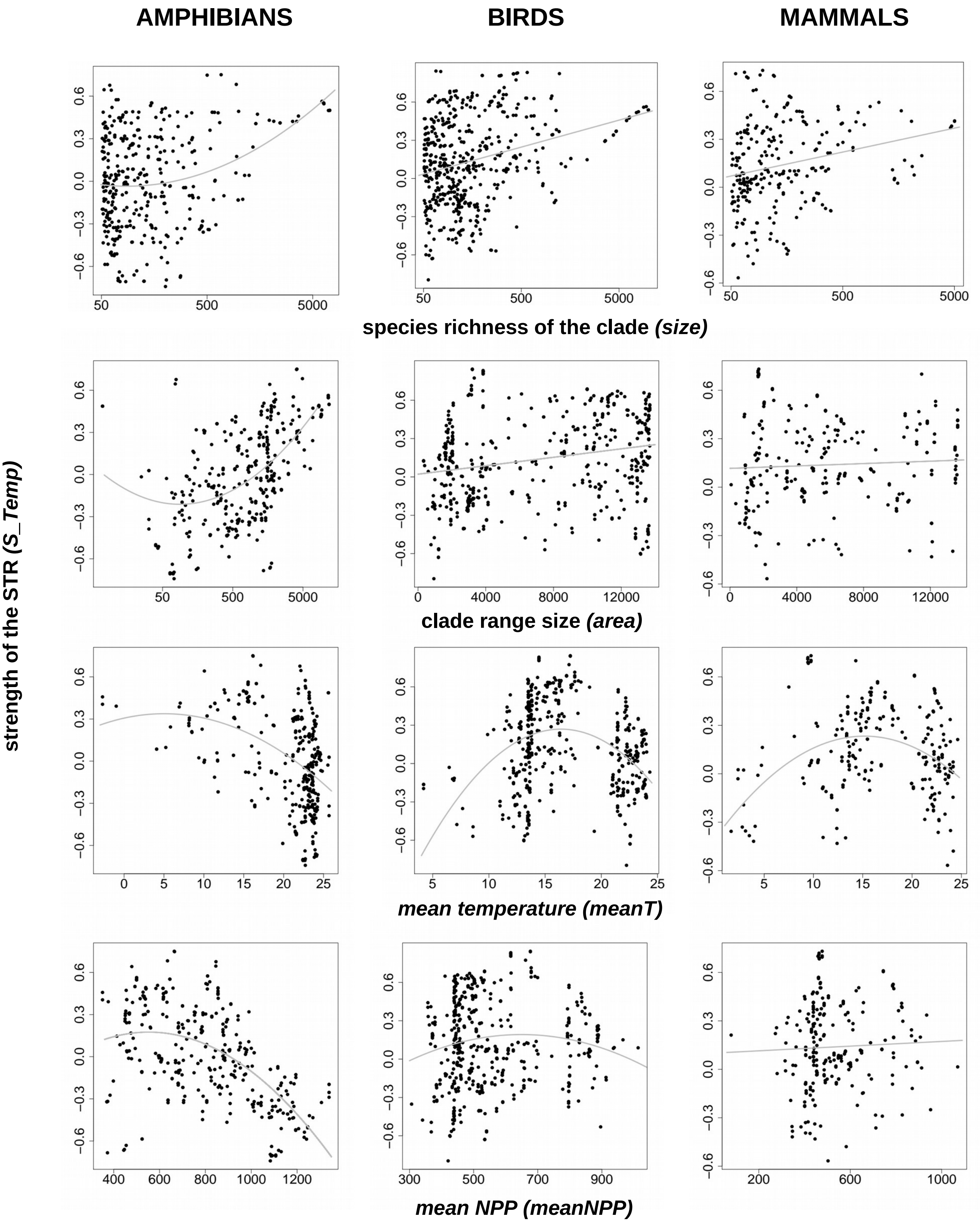

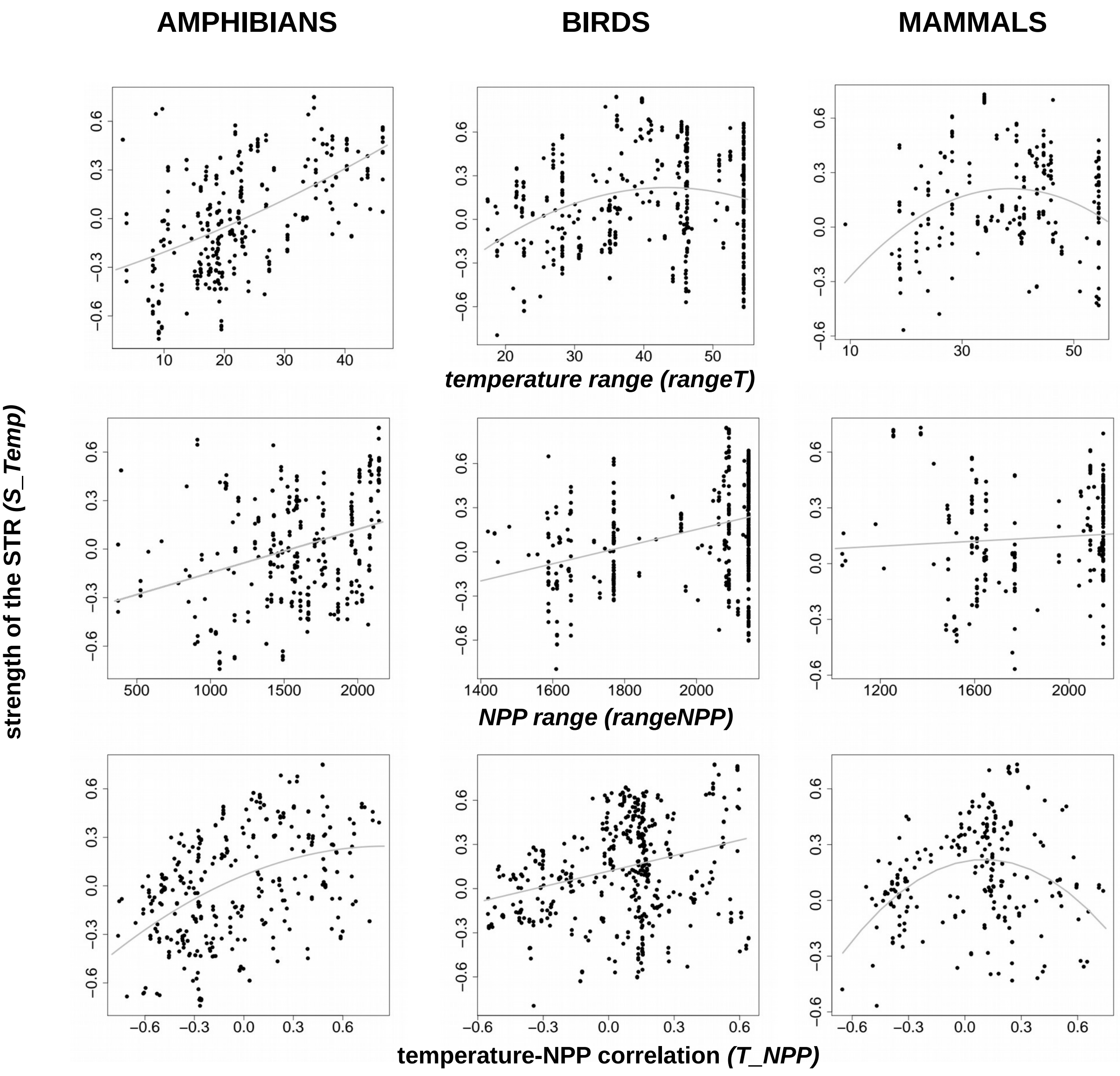
AMPHIBIANS
BIRDS
MAMMALS
strength of the STR (S_Temp)
temperature range (rangeT)
NPP range (rangeNPP)
temperature-NPP correlation (T_NPP)

**Figure S3.** The relationships between the SPR strength (*S_NPP*) and each explanatory variable (rows) for CLADES (all the PGLS univariate models are significant, see Table 2b). Grey lines represents univariate models in Table 2b, detailed description of the variables is in Table 1.

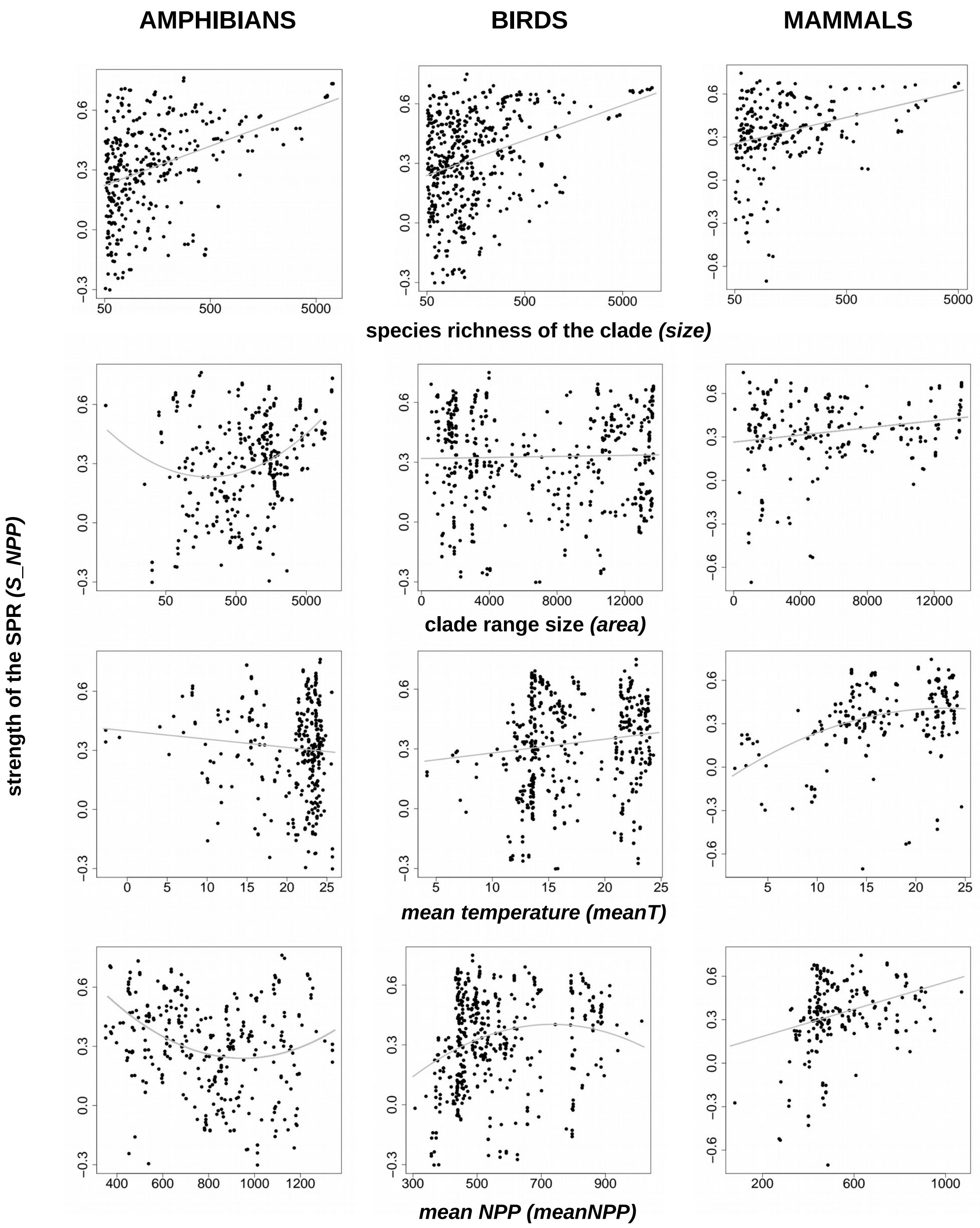

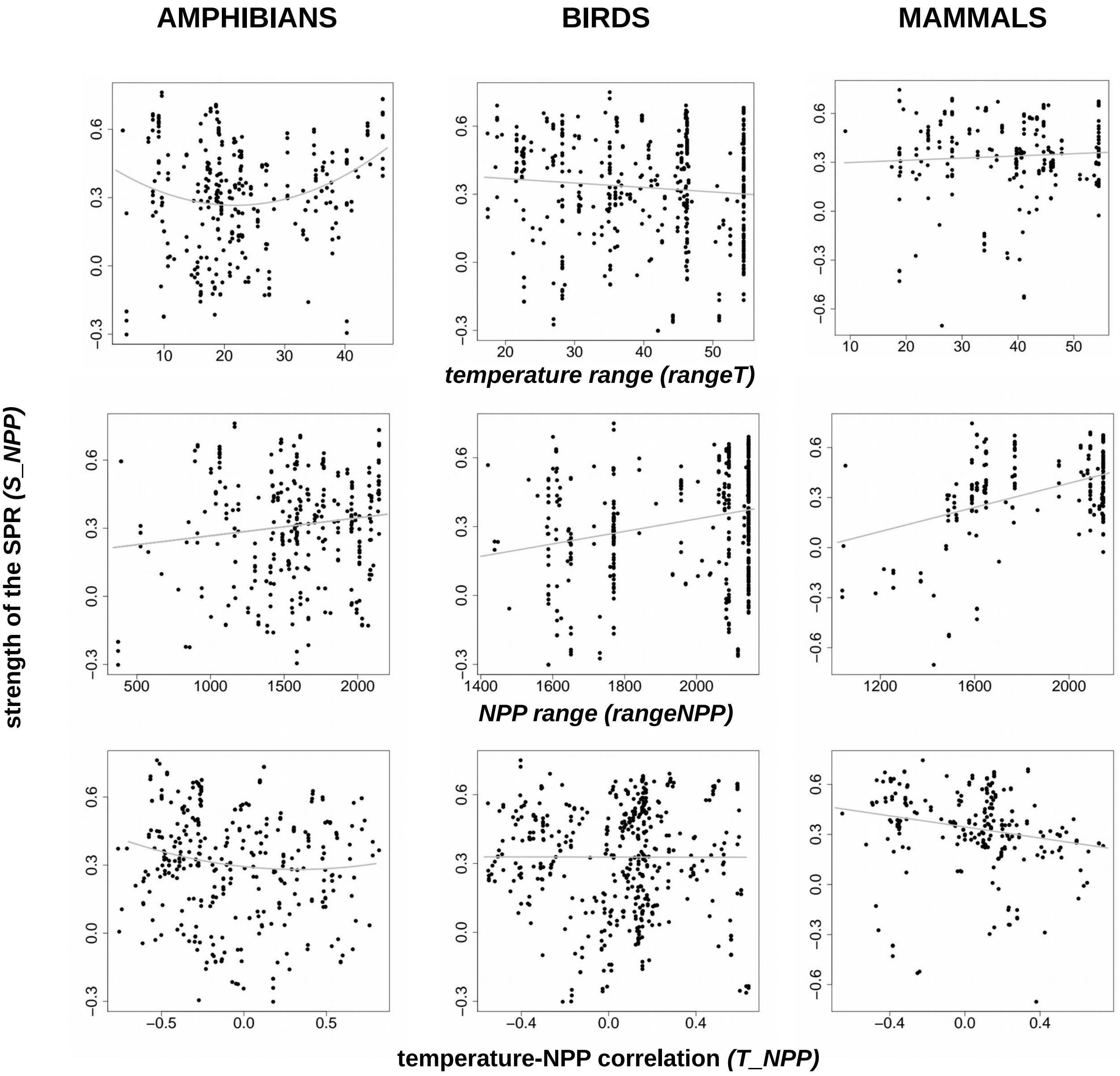

AMPHIBIANS
BIRDS
MAMMALS
strength of the SPR (S_NPP)
temperature range (rangeT)
NPP range (rangeNPP)
temperature-NPP correlation (T_NPP)

SUPPORTING ANALYSIS

As we argue in the main text, most STR and SPR relationships are monotonic, so the Spearman's correlation coefficient is a proper measure of their strength. Nevertheless, here we provide analysis using the coefficient of determination (R-squared, $R^2$) of the OLS model (either linear or quadratic). The rationale is that R-squared could potentially better express the strength of the relationship for hump-shaped or U-shaped relationships.

Specifically, for each dataset (for REGIONS) and each clade (for CLADES) we performed OLS regression model explaining (log) number of species by temperature or NPP, respectively. We performed these models in both linear and quadratic form and then chose the model better explaining variation in species richness using AIC comparison (if the AIC difference was lower than 2, we considered the relationship linear, otherwise we used the model with lower AIC). As we show in the main text (Fig. 2), some relationships (especially STRs) are negative. To make the results comparable to the results that used Spearman's correlation coefficient, we added plus or minus sign to the R-squared values, to indicate the direction of a relationship. The sign was determined based on the slope (positive or negative) of the linear OLS regression model.

Fig. S4 shows that both measures of STRs and SPRs strength are well correlated. This indicates that most of the relationships are indeed monotonic - U-shaped and hump-shaped relationships would not be well characterized by Spearman's correlation coefficient (being close to zero), and the correlation coefficient then would not correlate with the R-squared. It means that (1) the Spearman's correlation coefficient is a good measure of the STR and SPR strength and (2) it is reasonable to add plus or minus signs to the R-squared values indicating the direction of a relationship (which would not be the case for truly hump-shaped and U-shaped relationships).

Using this R-squared values as the response variable we performed the same analyses as for the correlation coefficient (see the description in the Material and methods section in the main text). Here, we present the results for the full models, univariate models, and variation partitioning. In REGIONS, we were able to explain 52% of variation in the strength of STRs (*S_Temp.R2*) and 19% of variation in the strength of SPRs (*S_NPP.R2*) by our explanatory variables (adjusted $R^2$ of the WLS full model). In CLADES, the full models explained 30% of the variation in STRs strength and 28% of the variation in SPRs strength for amphibians, 22% of the variation in STRs strength and 13% of the variation in SPRs strength for birds, and 22% of the variation in STRs strength and 19% of the variation in SPRs strength for mammals (adjusted $R^2$ of the PGLS full models). The results of the univariate models are presented in Table S2 and the results of the variation partitioning in Fig. S5.

**Figure S4**. The relationships between two measures of (a) the STR strength and (b) the SPR strength: R-squared of the OLS model, used in the supporting analysis (x-axis), and Spearman's correlation coefficient, used in the main text (y-axis). The first row comprises datasets of various ectothermic taxa (REGIONS), following rows comprise vertebrate clades (within amphibians, birds and mammals; CLADES).

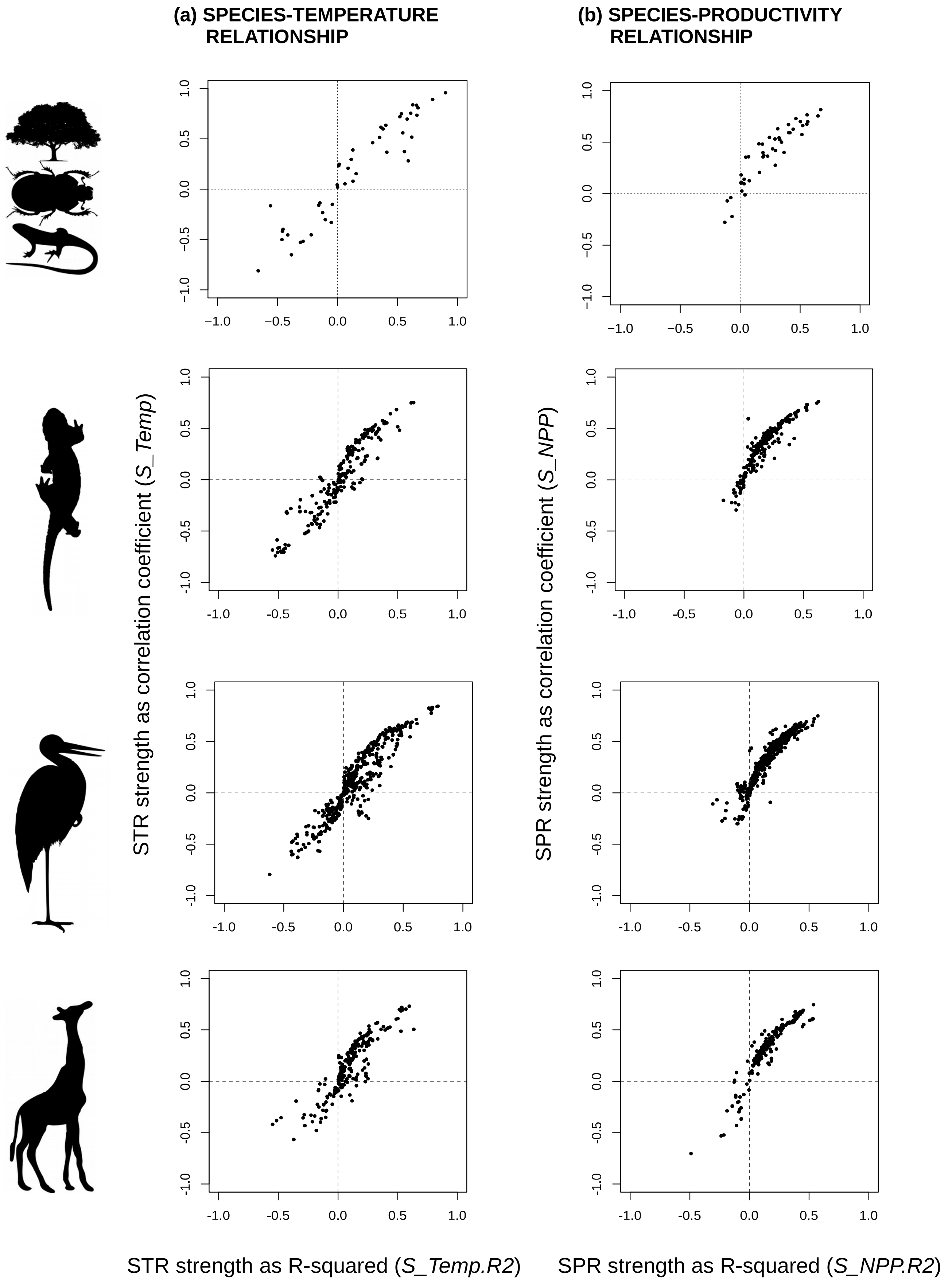

**Table S2**. Results of the univariate models explaining the variation in the strength of the STR (left column) and in the strength of the SPR (right column) expressed as R-squared for (a) datasets of various ectothermic taxa – REGIONS, and for (b) vertebrate clades – CLADES. The effects significant on α=0.05 are in bold, for the explanation of the variables see Table 1. AIC corrected for small sample size (AICc), adjusted $R^2$ (adj. $R^2$) and coefficients (coefs.) are displayed.

| | SPECIES-TEMPERATURE RELATIONSHIP | | | | SPECIES-PRODUCTIVITY RELATIONSHIP | | | |
|---|---|---|---|---|---|---|---|---|
| (a) | **Model** | **AICc** | **adj. $R^2$** | **coefs.** | **Model** | **AICc** | **adj. $R^2$** | **coefs.** |
| REGIONS | ***meanT*** | **35.2** | **0.53** | - | ***meanT*** | **-8.7** | **0.18** | + |
| | ***T_NPP*** | **44.3** | **0.43** | + | ***T_NPP+T_NPP***$^2$ | **-5.5** | **0.15** | -;+ |
| | ***rangeT + rangeT***$^2$ | **56.3** | **0.28** | -;+ | log(*size*) | -2.2 | 0.06 | + |
| | log(*area*) | 63.7 | 0.00 | + | *rangeT* | -0.4 | 0.02 | - |
| | log(*size*) | 64.8 | 0.00 | - | *log(grain)* | 1.1 | 0.00 | - |
| | *rangeNPP* | 65.0 | 0.10 | - | *rangeNPP* | 1.3 | 0.00 | + |
| | log(*grain*) | 66.8 | 0.00 | - | *meanNPP* | 1.5 | 0.00 | - |
| | *meanNPP* | 69.4 | 0.01 | - | *log(area)* | 1.6 | 0.00 | + |
| (b) | | | | | | | | |
| AMPHIBIANS | ***T_NPP+T_NPP***$^2$ | **-429.7** | **0.23** | +;- | **log(*size*)** | **-577.9** | **0.17** | + |
| | ***meanNPP*** | **-416.1** | **0.20** | - | **log(*area*)+log(*area*)**$^2$ | **-548.5** | **0.10** | -;+ |
| | **log(*area*)+log(*area*)**$^2$ | **-416.0** | **0.20** | -;+ | ***meanNPP*** | **-545.0** | **0.09** | - |
| | ***meanT+meanT***$^2$ | **-414.8** | **0.20** | +;- | ***meanT*** | **-542.1** | **0.08** | - |
| | ***rangeT+rangeT2*** | **-405.6** | **0.18** | -;+ | ***rangeT+rangeT***$^2$ | **-539.2** | **0.08** | -;+ |
| | **log(*size*)+log(*size*)**$^2$ | **-367.4** | **0.08** | -;+ | ***T_NPP+T_NPP***$^2$ | **-529.3** | **0.05** | +;+ |
| | ***rangeNPP*** | **-347.6** | **0.03** | + | ***rangeNPP*** | **-527.4** | **0.04** | + |
| BIRDS | ***meanT+meanT***$^2$ | **-516.7** | **0.14** | +;- | **log(*size*)** | **-881.1** | **0.09** | + |
| | ***T_NPP*** | **-516.1** | **0.14** | + | ***meanNPP+meanNPP***$^2$ | **-843.0** | **0.02** | +;- |
| | **log(*size*)** | **-502.8** | **0.11** | + | *T_NPP+T_NPP*$^2$ | -835.6 | 0.01 | -;+ |
| | ***rangeT+rangeT***$^2$ | **-492.6** | **0.10** | +;- | *area* | -835.5 | 0.01 | + |
| | ***meanNPP+meanNPP***$^2$ | **-491.1** | **0.06** | +;- | *rangeNPP* | -835.5 | 0.01 | + |
| | ***area*** | **-453.3** | **0.02** | + | *rangeT* | -833.6 | 0.00 | + |
| | ***rangeNPP*** | **-449.3** | **0.01** | + | *meanT* | -831.9 | 0.00 | - |
| MAMMALS | ***meanT+meanT***$^2$ | **-353.4** | **0.16** | +;- | **log(*size*)** | **-412.4** | **0.09** | + |
| | ***log(size)*** | **-345.9** | **0.12** | + | ***meanT+meanT***$^2$ | **-399.7** | **0.05** | +;- |
| | ***T_NPP+T_NPP***$^2$ | **-343.7** | **0.12** | +;- | *rangeT* | -391.4 | 0.01 | - |
| | ***area*** | **-322.4** | **0.04** | + | *area* | -391.3 | 0.01 | + |
| | *rangeT* | -317.1 | 0.02 | - | *T_NPP* | -389.2 | 0.00 | - |
| | *rangeNPP* | -316.6 | 0.02 | + | *rangeNPP* | -288.8 | 0.00 | + |
| | *meanNPP* | -313.3 | 0.00 | + | *meanNPP* | -388.8 | 0.00 | + |

**Figure S5**. Venn diagrams describing variation partitioning for (a) the strength of the STR expressed as R-squared (*S_Temp.R2*) and (b) the strength of the SPR expressed as R-squared (*S_NPP.R2*) for different data (rows). Within each panel, in the left column, explanatory variables are divided into 3 categories - phylogenetic scale (variable *size*), spatial scale (variables *area* and *grain* [*grain* only for REGIONS]) and region characteristics (three region characteristics with the strongest univariate effects on *S_Temp.R2* or *S_NPP.R2*, respectively, see Table S2 and Table 1). In the right column, the variation partitioning is further performed for these three region characteristics. The numbers represents adjusted $R^2$ (in %), zero and negative values are not displayed. Every section is coloured according to its adjusted $R^2$ value on the white-black scale (white corresponds to 0, black corresponds to 50). *Note*: *T_NPP* abbreviates the correlation between temperature and NPP.

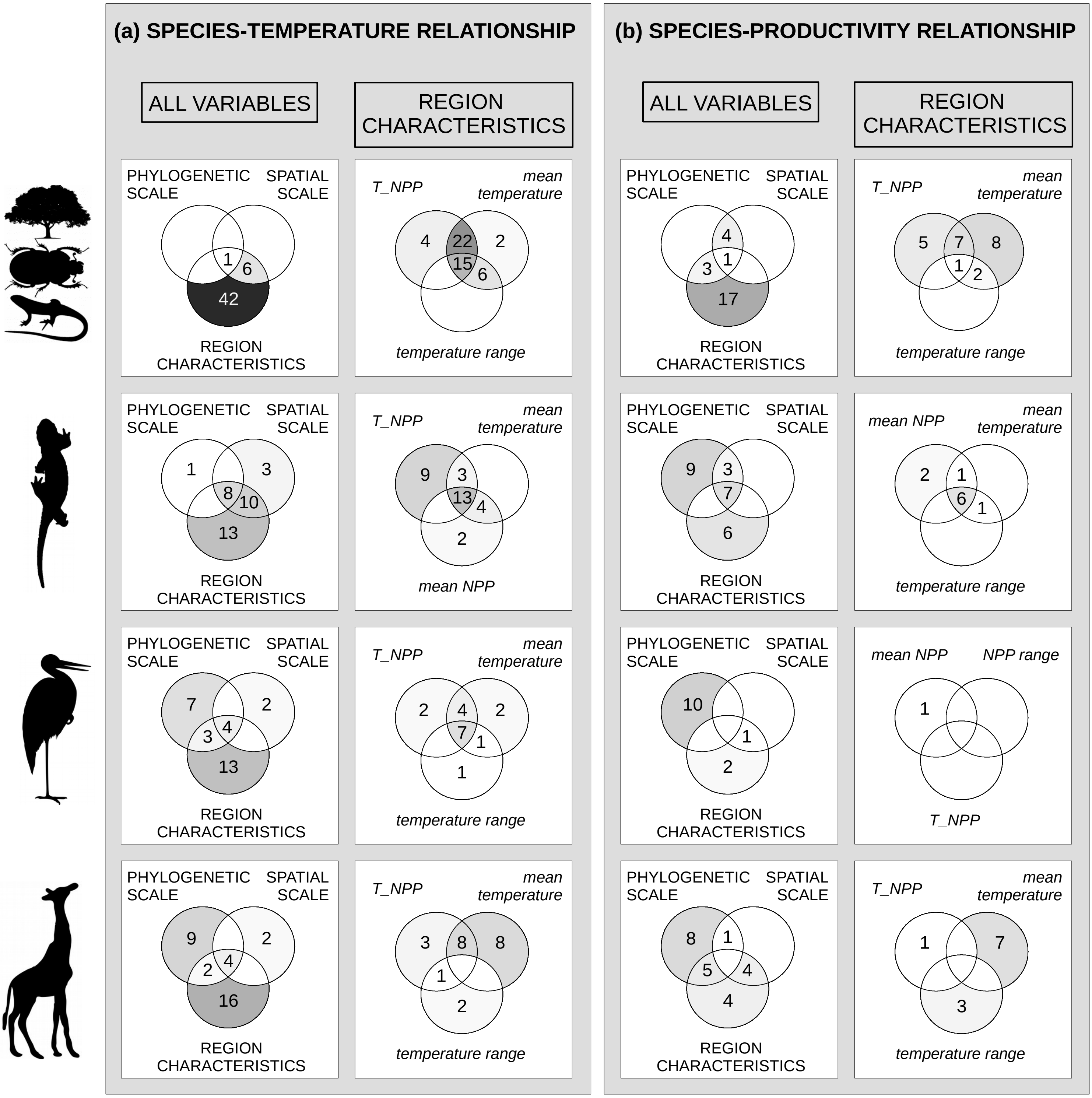
